# Perception/action coupling is modulated by the age-related motor skills of the agent performing the action

**DOI:** 10.1101/2024.08.24.609171

**Authors:** Nicole Clavaud-Seon, Quentin Guillon, Christina Schmitz

## Abstract

The perception/action coupling, underpinned by Mirror Neurons, allows the understanding of others’ action goal and intention picked up from cues conveyed by the individual’s kinematics and the context of the action. This mechanism can be modulated by familiarity with the observed action, by motor experience and motor expertise for that action, but could it be modulated by the age-related motor skills of the agent performing the action? We used an eye-tracking visual preference paradigm to study the modulation of perception/action coupling in 62 adults when viewing videos of daily actions presented in a forward reading direction or, for the same action, in a backward reading direction. Video actions were performed by young actors aged 4, 8, and 13 years and by adults. We found greater pupil dilation for all backward videos compared to forward videos. Interestingly, pupil dilation was greater for the forward videos with child actors than for those with adolescent/adult actors. An overall comparison of looking times during the preference phase did not reveal a preference for either video but testing for a context effect revealed a preference for backward videos when the last video observed in the exposure phase was forward. This study demonstrated the influence of the agent’s age-related motor abilities on the observer’s perception/action coupling and revealed that pupil dilation and the context effect could be relevant cues for exploring this coupling in a non-invasive, passive experimental setup that is particularly suitable for exploring perception/action coupling in very young children with a developmental disorder.

## 1. Introduction

We spend our time interacting with our environment, using our body and brain as tools to perform everyday actions, to observe and learn new ones mostly by imitation. But we do not suspect how complex it might be to perform even a nearly automatic daily action, because it seems so simple to us as adults. Indeed, an action can be considered as a coherent set of well-planned movements units oriented towards a goal, with varying degrees of finesse and complexity in the kinematics and with precise biomechanics that give coherence to the all from the beginning to the end. More precisely, what produces coherence to the perceived action is also the velocity in the execution of the movement, the grip strength, acceleration and jerk that are part of the kinematics. Motor learning and motor experience shape action since an early age and throughout life span.

Performing an action requires a set of neural mechanisms that are also involved when we try to understand an action performed by someone else. This mechanism, which enables us to couple the observed action with the motor representations in memory through motor simulation, is underpinned by mirror neurons. The Mirror Neuron Mechanism (MNM) (Rizzolatti et al., 2009) is also known as perception/action coupling and would allow a fine and immediate understanding of non-verbal gestures, the goal of others actions and their intentions also during social interaction observation. The MNM seems to be already functioning at an early age (Gallese et al., 2009) and would be involved in imitation, empathy, theory of mind and language. More specifically, predicting the goal of an action and the intention of the individual performing it would require inferences from the cues conveyed by the individual’s behavior and kinematics, as well as the context in which the action takes place (Iacoboni et al., 2005; Decroix & Kalenin, 2019; Stapel et al., 2012). Many factors can modulate the perception/action coupling (Rizzolatti & Fabbri-Destro, 2010) such as familiarity with the observed action (Calvo-Merino et al., 2005; Cross et al., 2006), motor experience (Van Elk et al., 2008) and even motor expertise for that action (Calvo-Merino et al., 2006).

Another factor that would modulate the perception/action coupling of an observer is the kinematics of the agent performing the action. When observing a child performing an action, for instance catching a ball, we get the impression that she or he is doing it in a very similar way to an adult. Nevertheless, motor development follows a long process of maturation from birth to adolescence that involves complex and interdependent cognitive and sensorimotor mechanisms such as planning of the movement, coordination between the limbs, execution and control of the gesture (Ripoll et al., 1994). As a consequence, action representations mature throughout childhood and adolescence with pivotal periods of motor development at 6/7 years and adolescence (Assaiante, 2012). This suggests that children’s gross and fine motor skills are different from those of adults, which also means that the kinematics of movements differ, among others, in finesse and precision (Kuhtz-Buschbeck et al., 1998; Pangelinan et al., 2010). Therefore, although actions performed by children seem to be similar to those of adults, their kinematics may differ. The more similar our kinematics is to the observed model, the greater the motor resonance (Cook et al., 2016), and the better the visual recognition performance (Casil and Giese, 2006). Hence, when we watch someone perform an action, could our perception/action coupling be modulated by the age-related motor skills of the agent performing the action?

To answer this question, we used an ecological, passive and non-invasive eye-tracking visual preference paradigm presenting with videos of daily actions (Clavaud et al., submitted). This paradigm allowed us to explore the perception/action coupling in children watching videos of adult actors performing daily actions, through the exploration of visual exploration behaviour cues such as looking times and pupil diameter variations over time. The aim of the current study was to investigate the modulation of perception/action coupling in adults with completed motor skills when viewing videos of daily actions performed by young actors aged 4, 8, and 13 years and by adults. We therefore sought to test the effect of a level of motor development varying according to the age of the actors. These actions presented with a variable perception/action coupling depending on whether a video was presented in the forward reading direction (i.e. allowing a strong coupling between the perceived action and the action the participant can perform), or the same video presented in the backward reading direction (in this case, the coupling would be weaker). We hypothesized that the more the actor’s motor skills differed in terms of motor development from those of the observer, the more a violation of the observer’s expectations would be measured, reflecting a difference in perception/action coupling. Following a previous study in children (Clavaud et al., submitted), we expected, in general, greater pupil dilation for backward actions in the exposure phase of our paradigm. Moreover, we expected this pupillary dilation to be all the more important as the actor was young, highlighting the impact of a different and immature kinematics on the matching between the perceived action and the representation of this action. In the preference phase an increase in looking time toward backward actions relative to forward actions was hypothesized, also greater for actions performed by young actors.

## 2. Material and methods

### 2.1. Participants

In total, 62 adults (33 females, 29 males) between the age of 22 and 60 years with mean age score of 32,9 (SD = 9,9) were included in this study. The experimental protocol was approved by the local ethics committee CPP Sud Est 1 N° ID-RCB : 2019-A01864-53. Participants were adults, with no neurological or psychiatric disorders, no seizures, no oculomotor disorders or visual exploration disorders and had to have normal to corrected vision with lenses or glasses. The absence of motor impairments or dyspraxic disorders in childhood was confirmed by means of a questionnaire in which the acquisition of the main motor milestones was assessed. All participants fully consented to be part of the study, and received a small gift in appreciation of their contribution.

### 2.2. Stimuli

252 videos of daily actions (as lace up a shoe, fold a T-shirt or put on a jacket) were created. Videos were shot in a neutral room with constant ambient light set at 100 lux. Four pairs of actors and actresses aged 4, 8, 13 and adult, dressed in neutral clothing, were filmed while performing the actions. They were filmed from a three-quarters point of view so that the actions were clearly visible to the observers. The start and end positions of the actors were similar and also kept constant across actions so that regardless of the direction of the video (played forward or backward), the start and end positions were the same. Videos were filmed in black and white with a Canon EOS 750 D digital camera. Then, with the iMovie’9 (version 10.1.8) software, the sound of the videos was suppressed and the reading direction of the videos was reversed so as to create, from the same daily action video, two videos: one video in forward reading direction and a second video in backward reading direction. Finally, they were set to 540×720 pixels in resolution. Thus, for the daily action « to put on a jacket », a first video was recorded in the forward direction (hereinafter referred to as “Forward”) and a second video was recorded in the backward direction (hereinafter referred to as “Backward”) and became the action « to take off a jacket ». Each video presented a plausible action, whether it was presented in the forward or backward direction. Finally, each forward and backward video was then doubled using Avidemux software (version 2.7.0), so that for the same action there were two “mirror” videos, one where the actor performing the action was positioned on the right side and the other where the actor was positioned on the left side. From these videos, 4 sets were created, each containing 7 blocks of 9 videos. Each participant viewed only one of the four randomly selected sessions. Videos of each set were randomised across participants. The order of appearance of backward and forward as well as the side of appearance of left or right videos were counterbalanced between trials.

### 2.3. Eye tracking

Pupil size and eye movements were measured using Tobii Pro X3-120 eye-tracker (Tobii Pro AB, Stockholm, Swedeen) in a free-head mode and at a sampling rate of 120 Hz (binocular). Participants sat in a comfortable armchair at a distance of 60 cm from a 20’’ display screen (1280×1024 pixels). The experiment took place in a soundproofed room with indirect lighting kept constant across participants (illuminance level ∼ 10 lux). The experiment was implemented using E-Prime 3 (Psychology Software Tools, Inc, Sharpsburg, USA).

### 2.4. Procedure

After the participant was comfortably seated, a photomotor reflex measured the diameter of each pupil at its minimum and maximum dilation. The photomotor reflex procedure consisted of a succession of black screen (2s), white screen (2s) and finally black screen (2s). The experimenter then presented as an example a couple of videos displaying an action in the forward and in the backward direction, and a five-point calibration was performed to ensure the good quality of the data. Then, the instruction given to each participant was to watch the videos of the daily actions carefully, as if they were at home quietly watching television. After each set, they were asked if they were able to distinguish between the two videos (backward and forward) for each action.

A trial consisted of an exposure phase followed by a visual preference phase (Figure 1). In the exposure phase, a grey screen with a central fixation point (and an associated sound) was first shown to participants for 500 ms. Next, either on the left or the right side of the screen, the first video of an action was presented for 6000 ms. Afterwards, the same sequence of events was repeated with the same action but played in the other direction and on the opposite side of the screen. In the preference phase, the test started only if the participant fixated the central fixation point on the grey screen. The same two videos were then displayed side by side on the screen, with each video remaining on the side where it appeared in the exposure phase.

**Figure 1.**
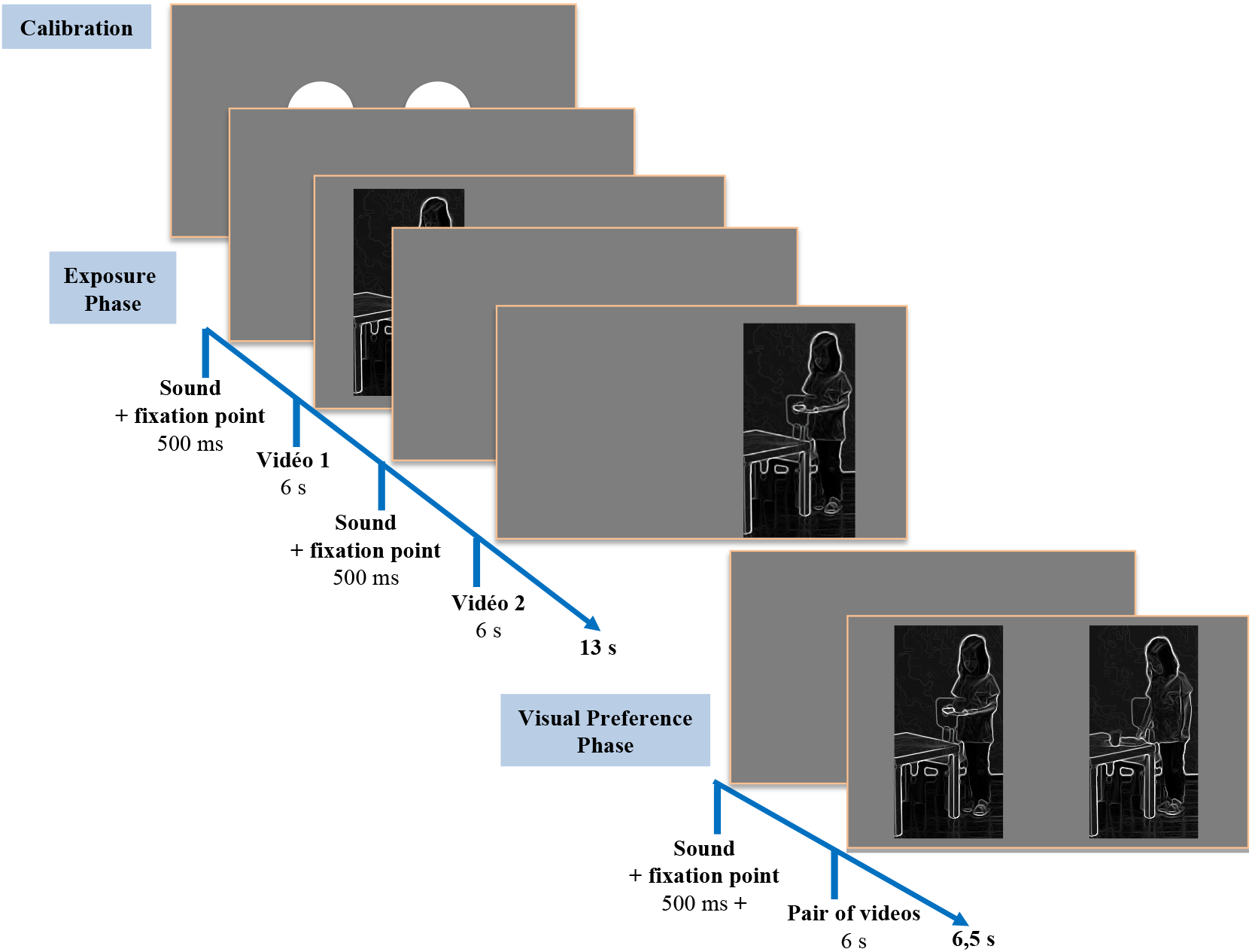
Experimental design showing the temporal sequence of the Exposure phase and the Visual preference phase.

### 2.5. Pre-processing of pupil size and eye movement data

All preprocessing stages were performed into Matlab (R2014). Only trials with less than 30% of missing data were analyzed, resulting in the removal of 107 (2.7%) trials out of a total of 3906. The median number of trials analyzed per participant is 62 (IQR = 62 - 62). One participant had only 26 trials (out of a total of 63) meeting this condition, and was excluded from the analyses.

Blinks and related artifacts (opening and closing periods of the eyelid) were first identified using an algorithm to distinguish between what is noise in the pupillary signal related to device measurement error and what is actually a blink and its artifacts (Hershman et al., 2018).

#### Pupil size

Blink and blink-related artifacts were first linearly interpolated. After interpolation, pupil data were smoothed using a zero-phase low-pass Butterworth filter (4Hz) to remove fast instrument noise, and down sampled from 120 to 40 Hz by taking the median pupil size per time in (25ms) to reduce computational cost. Pupil data were then aligned to the onset of each action video and segments of interest were created by extracting data between -100 pre- and +6225 ms post-video onset. Baseline correction (average pupil size in the time window from -100 to 100 ms) was applied using a subtractive method on a segment-by-segment basis. Segments where blinks and blink-related artefacts exceeded 30% of the total segment sample were rejected from further analysis. A total of 7541 segments were retained (out of 7812). The median number of segments per participant was 124 (IQR = 122 - 124). No participants were excluded. For each participant, the winsorized mean (\alpha = 0.1) of pupillary response over time was calculated for both forward and backward actions.

#### Eye movements

Total looking time towards either backward or forward actions during the exposition phase was computed by multiplying the total number of gaze points falling onto the respective action video by the sampling time interval (i.e. 8.333 ms). For the preference phase, preference for backward relative to forward actions was computed by dividing the number of gaze points that fell onto the backward action by the total number of gaze points that fell onto any actions (backward + forward). Thus, the preference represents the probability of looking at backward actions, given that the participant is looking at an action, whether backward or forward, and range from 0 to 1. Trials were excluded if 1) only one of the two actions was looked at (defined as less than 125 ms of looking time) exclusively, either backward (n=136) or forward (n=179) and/or 2) the percentage of blink or blink-related artefact samples exceed 30% during the preference phase (n=52). This last criteria was decided upon inspection of the distribution of the number of missing gaze data points across the whole sample. As a result, a total of 3504 trials (90%) were analysed (out of 3906). The median numbers of trials per participants out of 63 was 59 (IQR = 55 - 61). No participants were excluded from these analyses.

### 2.6. Statistical analyses

We used R (Version 3.6.1; R Core Team 2019) and the R-packages *lme4* (Bates et al., 2015), *lmerTest* (Kuznetsova et al., 2017), *tidyverse* (Wickham et al., 2019), *WRS2* (Mair & Wilcox, 2020) for data preparation, analysis and presentation.

#### Curve shape of pupil change

We performed a growth curve analysis of the pupil size (Mirman et al., 2008). Growth curve analysis is a multi-level modelling technique that allows the changes in pupil size to be modelled with orthogonal polynomials representing key aspects of the curve shape. For the current study, the pupil time course was modelled with a third-order polynomial and fixed effects of direction (backward vs. forward), the age of the actor performing the action (aged 4, 8, 13 years and adults) and their interaction on all polynomials. To simplify the analyses and, we grouped together the videos performed by the younger actors (4 and 8 years of age, hereinafter referred to children) and the videos performed by the older actors (13 years and adults, hereinafter referred to ado/adults). In this model, the intercept refers to the mean pupil size over the full segment; the linear term refers to the slope of the pupil time course, with lager values indicating lager pupil size at the end than at the beginning of the segment; the quadratic term refers to the shape of primary inflection, with more positive values indicating a flatter inverted-U shape and more negative values, a steeper inverted-U shape (Kuchinsky et al., 2013). The random structure included participant random effects and participant-by-condition random effects on all time terms (i.e. intercept varying among participants and among directions and ages of the actors within participants). The models were fitted with maximum likelihood estimation and parameter estimates, degrees of freedom and corresponding p-values were estimated using the Satterthwaite method.

### Gaze behaviour

Given the bounded nature of the preference score (between 0 and 1), generalized linear mixed effects models (GLMMs) were used. The GLMMs were fitted with maximum likelihood estimation using a binomial distribution and a logit link function. The random structure included random effects per observation (OLRE procedure) to take into account the over dispersion of residuals usually observed with this type of data. Further, to control for the context effect that may be induced by the presentation order of the forward and backward videos during the exposure phase, we tested whether the total looking time measured during the visual preference phase was influenced by the last action video viewed during the exposure phase. The context of presentation of the videos during the exposure phase, the age of the actors performing the action and their interaction were considered as fixed effects.

Odds ratios (ORs), and their 95% confidence intervals calculated by bootstrapping, are reported to facilitate interpretation and indicate the presence of a preference for backward (OR > 1) or forward (OR < 1) actions. A model comparison was carried out to test the statistical significance of the fixed effects of action order and actor age by comparing the change in residual deviance between the full model and a model without the target fixed effect using a likelihood ratio test.

## 3. Results

### 3.1. Pupil response

A significant interaction between the linear term, the direction of the actions and the age of the actors was found (Est = 0.12, ES = 0.05, p = 0.009), indicating that the effect on the linear term of the age of the actors was different depending on whether the actions were in the backward or forward direction. Similarly, a significant interaction was revealed between the quadratic term, direction of actions, and age of the actors, (Est = 0.08, ES = 0.03, p = 0.008), indicating that the effect on the quadratic term of the actor age was different depending on whether the actions were in the backward or forward directions (Table 1).

**Table 1.**

|  | *Est.* | *ES* | *df* | *t-value* | *p-value* |
| --- | --- | --- | --- | --- | --- |
| ***Fixed Effects*** |  |  |  |  |  |
| Intercept | 0.24 | 0.01 | 62 | 16.3 | < 0.001 |
| Linear term | 0.70 | 0.07 | 62 | 9.8 | < 0.001 |
| Quadratic term | -0.67 | 0.04 | 62 | -15.6 | < 0.001 |
| Direction (backward) | 0.02 | 0.00 | 62 | 4.9 | < 0.001 |
| Actors age (adolescents/adults) | -0.01 | 0.00 | 62 | -1.2 | 0.225 |
| Linear term x direction (backward) | 0.05 | 0.05 | 62 | 1.1 | 0.298 |
| Quadratic term x direction (backward) | -0.08 | 0.03 | 62 | -2.9 | 0.006 |
| Linear term x actors age (adolescents/adults) | -0.15 | 0.04 | 62 | -3.8 | < 0.001 |
| Quadratic term x actors age (adolescents/adults) | -0.18 | 0.03 | 62 | -6.9 | < 0.001 |
| Linear term x direction (backward) x actors age<br>(adolescents/adults) | 0.12 | 0.05 | 62 | 2.7 | 0.009 |
| Quadratic term x direction (backward) x actors age<br>(adolescents/adults) | 0.08 | 0.03 | 62 | 2.7 | 0.008 |

To better understand these interactions, the simple effects of the direction and age of the actors were conducted, allowing exploring the effect of one of these two variables on the linear and quadratic terms at each level of the other variable.

Concerning, the simple effect of the direction of the action (Figure 2), a significant interaction was found between the quadratic term and the direction in the condition where the actors performing the action were children (Est = -0.11, ES = 0.03, p < 0.001). Thus, when the actors were children the shape of the first inflection of the pupillary response was steeper for actions in the backward direction. When the actors were adolescents/adults, a significant interaction was found between the linear term and direction (Est = 0.15, ES = 0.05, p = 0.002), indicating that the slope of pupillary dilation was steeper in the backward direction than in the forward direction (Table 1).

**Figure 2.**
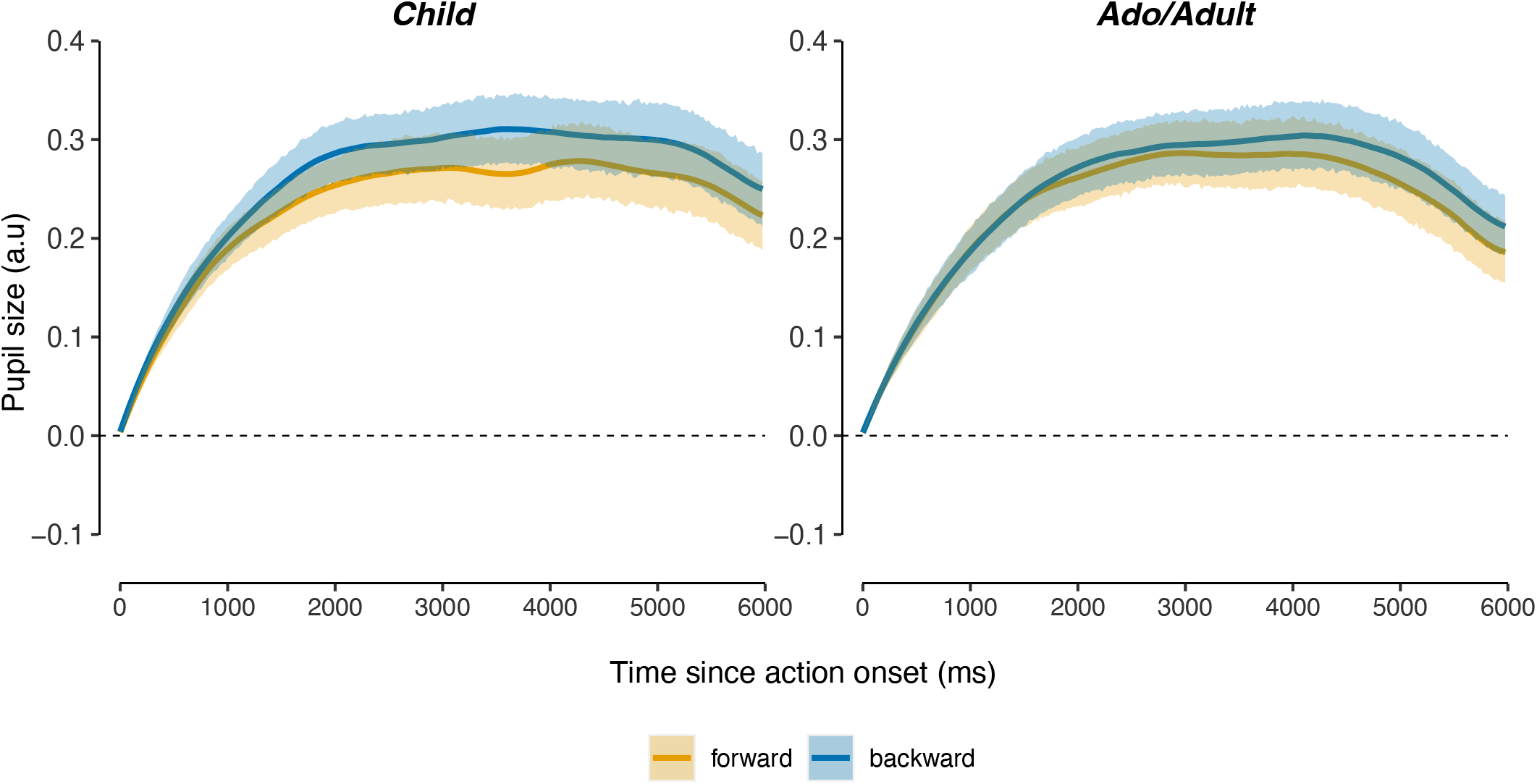
Time course of pupil size when the actors were children (left) or adolescents/adults (right), depending on the direction of the videos: Forward (orange) and Backward (blue). Solid lines represent the observed pupil size and shaded areas the 95% bootstrapped confidence intervals.

Concerning the simple effect of the age of the actors (Figure 3), a significant interaction was found between the quadratic term and the age of the actors when the actions were in the backward direction (Est = - 0.08, ES = 0.03, p = 0.007), indicating that when the actors were adolescents/adults, the shape of the first inflection in the condition where the actions were in the backward direction was steeper than when the actors were children. When the actions were in the forward direction, an interaction was found between the linear term and the age of the actors (Est = -0.10, ES = 0.05, p = 0.022), and between the quadratic term and the age of the actors (Est = -0.20, ES = 0.03, p < 0.001). This indicated that when the actions were in the forward direction, the pupillary response was less pronounced and steeper when the actors were adolescents/adults than when the actors were children (Table 1).

**Figure 3.**
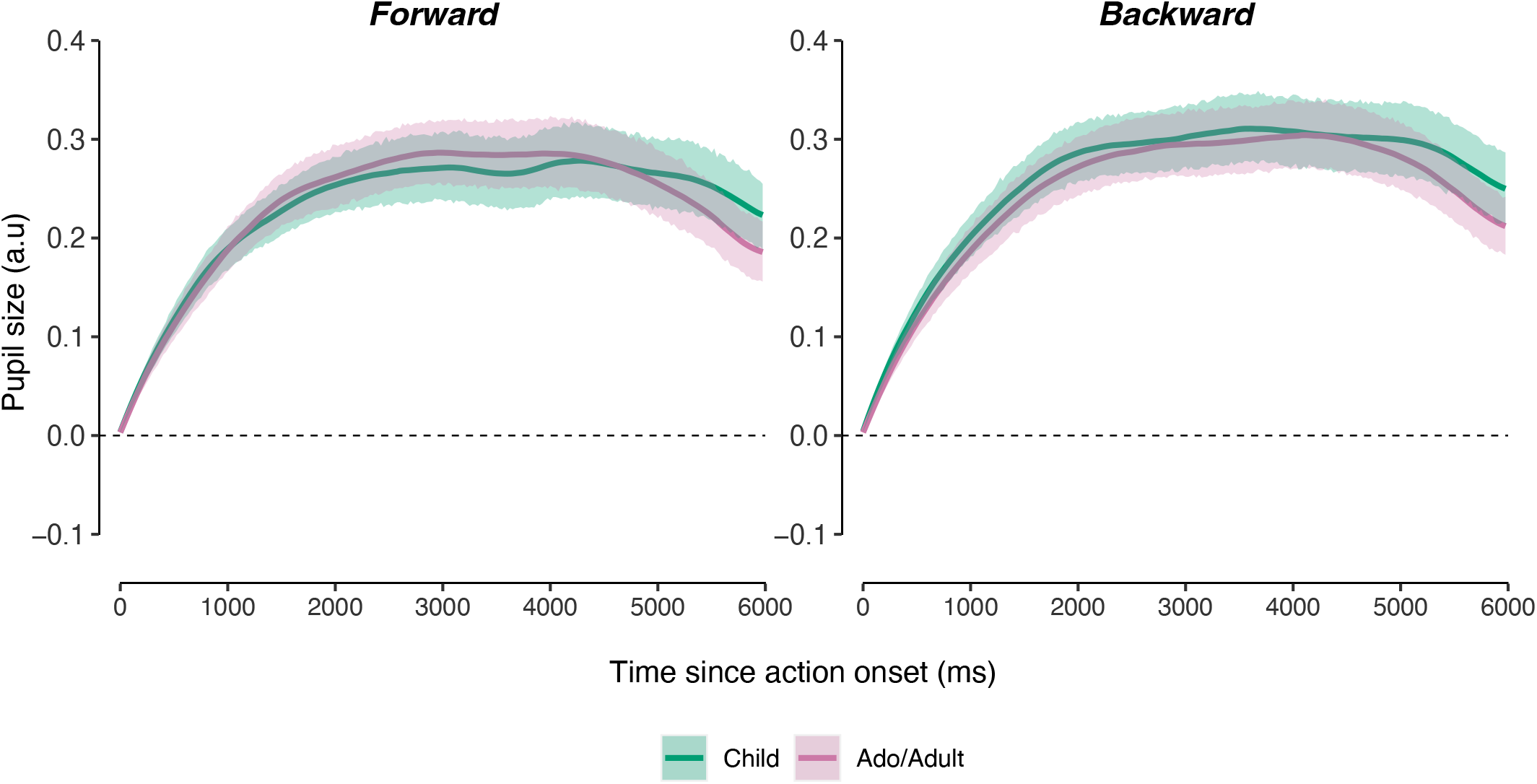
Time course of pupil size for Forward (left) and Backward (right) videos, depending on whether the actions were performed by a child (green) or an adolescent/adult (pink) actor. Solid lines represent the observed pupil size and shaded areas the 95% bootstrapped confidence intervals.

### 3.2. Preference

No simple effect of direction was found, as evidenced by no preference towards backward over forward videos. Indeed, accounting for the by-participants and by-actions variation, the odds of looking at backward videos relative to forward videos was 1.06 with a 95% bootstrapped confidence interval of [0.97, 1.15]. This indicated that all plausible values crossed the value 1.00 which indicates that the null hypothesis could not be rejected and that there was no preference for backward over forward videos.

A main effect of the context was found (X2(1)=76.11, p <0.001). In particular, a preference for backward videos was found when the video preceding the preference phase was forward (OR = 1.49, ICboot95%[1.34 - 1.67]). Conversely, a preference for forward videos was found when the video preceding the preference phase was backward (OR = 0.75, ICboot95%[0.65 - 0.85]). The main effect of the age of the actors was not significant (X2(1)=0.01, p = 0.923).

Finally, an interaction between context and age of the actors was found (X2(1)=5.79, p = 0.016), indicating a difference in the effect of context depending on whether the actors were children or a adolescents/adults. To better understand the origin of this interaction, the simple effects of context and age of the actors were examined.

Regarding the simple effect of the context of presentation, whatever the age of the actors, a context effect was revealed (X2(1)=38.02, p<0.001 and X2(1) =81.83, p<0.001), respectively for child and adolescent/adult actors. In the condition where the actors were children, a preference for backward videos was found when the video preceding the preference phase was forward (OR = 1.42, ICboot95% [1.28 - 1.60]) and conversely when the video preceding the preference phase was backward (OR = 0.80, ICboot95% [0.67 - 0.96]). Similarly, when the actors were adolescents/adults, a preference for backward videos was found when the video preceding the preference phase was forward (OR = 1.55, ICboot95% [1.37 - 1.79]) and conversely when the video preceding the preference phase was backward (OR = 0.71, ICboot95% [0.63 - 0.81]).

No significant difference was found on the simple effect of the age of the actors (X2(1)=2.89, p=0.089, and X2(1) =2.98, p=0.084, respectively for the forward and backward videos). When the video preceding the preference phase was backward, a preference for the forward video was found, independent of the age of the actors (OR = 0.80, ICboot95% [0.67 - 0.96; OR = 0.72, ICboot95% [0.64 - 0.83], for child and adolescent/adult actors respectively). Similarly, when the video preceding the preference phase was forward, a preference for the backward videos was found, independent of the age of the actors (OR = 1.42, ICboot95% [1.26 - 1.62]; OR = 1.55, ICboot95% [1.37 - 1.74], for child and adolescent/adult actors respectively).

Together, these results indicate that the context by age of the actor interaction indicates that the effect of the context was more pronounced when the actors were adolescents/adults.

## 4. Discussion

The main objective of this study was to characterize, via visual exploration and particularly pupil dilation and looking times, the spontaneous distinction of daily actions (presented in forward and backward directions) performed by actors who were either children aged of 4 and 8, or adolescents (aged 13) and adults. Consistent with our first hypothesis, we found greater pupil dilation for backward videos compared to forward videos. This effect was robust and found regardless of the age of the actors. Interestingly, we found greater pupil dilation for the forward videos when the actors were children, as compared to forward videos with adolescent/adult actors. An overall comparison of looking times during the preference phase did not reveal a preference for either video. Instead, testing for a context effect revealed a preference for backward videos exclusively when the last video observed in the exposure phase was forward.

### 4.1. The perception/action coupling is sensitive to the kinematics of the agent performing the action

In this study, the pupil dilation findings support our initial hypothesis that the perception/action coupling could be modulated by the agent’s age-related motor skills. We assumed this variation to be related to the disturbance of the kinematics of the actors’ movements only, rather than being related to an implausible action. Indeed, it could be argued that this effect could result from the fact that the reverse direction of the actions generates a quirk in the backward direction. Weirdness due, for example, to the violation of the laws of gravity exerted on objects, which could have made the videos backward more attractive and generate a novelty effect. However, we had controlled that all actions were plausible in both the forward and backward directions (e.g. “lace up a shoe” in the forward direction, becomes “unlace a shoe” in the backward direction) and we also controlled that no effect of gravity was visible on the objects. Thus, forward and backward videos could both correspond to the participant’s action representations. We suggest that greater pupil dilation for backward videos marked expectation violation and surprise (Lavin et al., 2014; Preuschoff et al., 2011), as the actors’ reversed kinematics differed from participants’ representations. Reversing the direction of the video resulted in a disruption of the kinematic dynamics (acceleration, deceleration, grip strength, jerk). Several studies have pointed out the importance of kinematics in the recognition and understanding of actions (Decroix & Kalenin, 2019; De Marco et al., 2020; Morita et al., 2012; Stapel et al., 2012; Troje et al., 2005). For example, kinematics modulations intrinsic to an observed reach-to-grasp movement are already informative about the motor intention of others (De Marco et al., 2020) and importantly, when motor information coming from an agent kinematics is not processed by the observer, the latter fails to identify the intention of the agent performing the action (Boria et al., 2019).

Further, we found an effect of the motor development of the actors for forward actions. Indeed, participants showed greater pupil dilation for forward videos performed by child actors compared to forward videos performed by adolescent/adult actors. This increase in pupillary response may reflect the surprise and uncertainty due to differences in the kinematics of the child actors, less precise, less fine, with more clumsiness and less fluidity, which may have been perceived by the adult participants, hence disrupting the perfect match between the observed action and their representation. Indeed, the differences in the kinematics of the 4- and 8-year-old actors underlie a different level of motor development, as compared to the participants who were adults with mature motor skills and long motor experience. This reinforces the idea that the closer one is to someone’s kinematics, the easier it is to perceive, predict, and interpret another’s actions (Cook, 2016; De Marco et al., 2020). This idea is all the more reinforced as we found a context effect of the video presentation in the exposure phase that impacted participants looking times in the preference phase. Indeed, when the last video viewed by the participants in the exposure phase was in the forward direction, in the preference phase the participants preferentially watched the backward video and conversely when the last video viewed was in the backward direction, in the visual preference phase the participants preferentially watched the forward video. This indicated that they differentiated well between the two actions, hence reflecting a good perception/action coupling. Interestingly, this context effect was significantly more important for videos with adolescent/adult actors, than for videos with child actors. This again may be related to the fact that the kinematics of adolescents/adults was more similar to those of adult participants due to closer motor skills. Hence, when the coupling was weaker, i.e. when watching videos of actions performed by child actors, the context effect was less marked as if dampen by uncertainty due to an imperfect matching with the participant’s representations.

### 4.2. Pupillary dilation and context effect could be better indexes of perception/action coupling than looking times

The results of this study confirm our hypotheses regarding pupillary dilation as a possible indicator of perception/action coupling, but question the validity of looking times as an index of this coupling. Indeed, we expected the violation of participants’ expectations to be revealed by greater looking times for backward videos in the preference phase, but there was statically no difference between the two conditions. Looking times include, among others, saccades and fixations and contribute to the orientation of explicit visuo-spatial attention (Jeannerod et al., 1968), which is a conscious process. During the preference phase, participants could have been influenced by the instruction given to them before the experiment. Indeed, after each set of 9 videos, they were asked if they could distinguish between the two directions of the videos (backward and forward) for each action. This may have induced a bias in their visual exploration behaviour during the preference phase, causing alternating exploration between the two action videos without clear preference. This active search might indeed have caused looking times not to be a reliable index of the perception/action coupling in our study.

To address this bias, we tested for the presence of a context effect. The paradigm design engaged participants to watch the videos as presented one after the other at first, and then to freely watch either of the two videos as they were presented simultaneously. If the context effect indicated an orientation of their attention towards the action opposite to the one they saw last in the exposure phase, this was modulated by the proximity with the aged-related motor skills of the actors performing the action. Hence, an attenuation of this context effect might sign uncertainty about the video that was just watched in the exposure phase, which makes it an interesting index.

On the contrary, participants cannot control pupillary dilation, which is a physiological unconscious response. Variations in pupillary diameter in response to a stimulus are related to the autonomic nervous system and precisely to the sympathetic activation (Vries et al., 2021 for review). Pupil variation is an involuntary marker of the autonomic nervous system activity, meaning it indexes its actual spontaneous activity, making pupillometry an objective measurement tool. According to Jackson and Sirois (2009), pupillometry is a more suitable and flexible method of measurement than the common Violation Of Expectation (VoE) method based on looking times that can be affected by order effects (Schöner & Thelen, 2006). Sirois and Jackson (2012) showed how pupil dilation is sensitive to perceptual dynamics in typical infant VoE studies, allowing for a much fine-grained interpretation of infant information processing, compared to looking time measures that these researchers consider more coarse-grained. Many studies have indeed demonstrated that pupillary response is a psychophysiological index that allows for finer observations and interpretations than looking times or eye movement alone (Morita et al., 2012; Sirois & Jackson, 2012), especially when looking times may lead to equivocal interpretations (Jackson & Sirois, 2009).

### 4.3. Conclusion

This study not only demonstrated the influence of the agent’s age-related motor abilities on the observer’s perception/action coupling, but also revealed that pupil dilation, as well as the context effect in an exposure/preference paradigm, could be relevant cues to this coupling. Further, the fact that experimental set-up is non-invasive, passive, and ecologically friendly make it particularly appropriate to explore the perception/action coupling in very young children, including those with developmental disorders.

## Acknowledgements

We are grateful to all participants, as well as the eight young models who kindly accepted to be filmed for the videos. We thank Lucie Gabard for her help with data acquisition.

## References

Assaiante, C. (2012). Action and representation of action during childhood and adolescence: a functional approach. Neurophysiologie Clinique/Clinical Neurophysiology, 42(1-2), 43–51.

Boria, S., Fabbri-Destro, M., Cattaneo, L., Sparaci, L., Sinigaglia, C., Santelli, E., Cossu, G., & Rizzolatti, G. (2009). Intention understanding in autism. PloS one, 4(5), e5596. 10.1371/journal.pone.0005596

Calvo-Merino, B., Glaser, D. E., Grèzes, J., Passingham, R. E., & Haggard, P. (2005). Action observation and acquired motor skills: An fMRI study with expert dancers. Cerebral Cortex, 15(8), 1243–1249. 10.1093/cercor/bhi007

Calvo-Merino, B., Grèzes, J., Glaser, D. E., Passingham, R. E., & Haggard, P. (2006). Seeing or Doing? Influence of Visual and Motor Familiarity in Action Observation. Current Biology, 16(19), 1905–1910. 10.1016/j.cub.2006.07.065

Casile, A., & Giese, M. A. (2006). Nonvisual motor training influences biological motion perception. Current Biology, 16(1), 69–74. 10.1016/j.cub.2005.10.071

Cook, J. (2016). From movement kinematics to social cognition: the case of autism. Philosophical Transactions of the Royal Society B: Biological Sciences, 371(1693), 20150372. 10.1098/rstb.2015.0372

Cook, J. L., Blakemore, S. J., & Press, C. (2013). Atypical basic movement kinematics in autism spectrum conditions. Brain, 136(9), 2816–2824. 10.1093/brain/awt208

Cross, E. S., Hamilton, A. F., & Grafton, S. T. (2006). Building a motor simulation de novo: observation of dance by dancers. NeuroImage, 31(3), 1257–1267. 10.1016/j.neuroimage.2006.01.033

Decroix, J., & Kalénine, S. (2019). What first drives visual attention during the recognition of object-directed actions? The role of kinematics and goal information. Attention, Perception, and Psychophysics. 10.3758/s13414-019-01784-7

De Marco, D., Scalona, E., Bazzini, M. C., Avanzini, P., & Fabbri-Destro, M. (2020). Observer-Agent Kinematic Similarity Facilitates Action Intention Decoding. Scientific reports, 10(1), 2605. 10.1038/s41598-020-59176-z

De Vries, L., Fouquaet, I., Boets, B., Naulaers, G., & Steyaert, J. (2021). Autism spectrum disorder and pupillometry: A systematic review and meta-analysis. Neuroscience and biobehavioral reviews, 120, 479–508. 10.1016/j.neubiorev.2020.09.032

Gallese, V., Rochat, M., Cossu, G., & Sinigaglia, C. (2009). Motor Cognition and Its Role in the Phylogeny and Ontogeny of Action Understanding. Developmental Psychology. 10.1037/a0014436

Hershman, R., Henik, A., & Cohen, N. (2018). A novel blink detection method based on pupillometry noise. Behavior Research Methods, 50(1), 107–114. 10.3758/s13428-017-1008-1

Iacoboni, M., Molnar-Szakacs, I., Gallese, V., Buccino, G., Mazziotta, J. C., & Rizzolatti, G. (2005). Grasping the intentions of others with one’s own mirror neuron system. PLoS biology, 3(3), e79. 10.1371/journal.pbio.0030079

Jackson, I., & Sirois, S. (2009). Infant cognition: going full factorial with pupil dilation. Developmental science, 12(4), 670–679. 10.1111/j.1467-7687.2008.00805.x

Jeannerod, M., Gerin, P., & Perrier, J. (1968). Deplacements et fixations du regard dans l’exploration libre d’une scene visuelle. Vision Research, 8(1), 81–97. 10.1016/0042-6989(68)90066-7

Krüger, M., Bartels, W., & Krist, H. (2020). Illuminating the Dark Ages: Pupil Dilation as a Measure of Expectancy Violation Across the Life Span. Child development, 91(6), 2221–2236. 10.1111/cdev.13354

Kuchinsky, S. E., Ahlstrom, J. B., Vaden, K. I., Cute, S. L., Humes, L. E., Dubno, J. R., & Eckert, M. A. (2013). Pupil size varies with word listening and response selection difficulty in older adults with hearing loss. Psychophysiology, 50(1), 23–34. 10.1111/j.1469-8986.2012.01477.x

Kuhtz-Buschbeck, J. P., Stolze, H., Jöhnk, K., Boczek-Funcke, A., & Illert, M. (1998). Development of prehension movements in children: a kinematic study. Experimental brain research, 122(4), 424–432.

Kuznetsova, A., Brockhoff, P. B., & Christensen, R. H. B. (2017). lmerTest Package: Tests in Linear Mixed Effects Models. Journal of Statistical Software, 82(13). 10.18637/jss.v082.i13

Lavín, C., San Martín, R., & Rosales Jubal, E. (2014). Pupil dilation signals uncertainty and surprise in a learning gambling task. Frontiers in behavioral neuroscience, 7, 218. 10.3389/fnbeh.2013.00218

Lawson, R. P., Mathys, C., & Rees, G. (2017). Adults with autism overestimate the volatility of the sensory environment. Nature neuroscience, 20(9), 1293–1299. 10.1038/nn.4615

Mair, P., & Wilcox, R. (2020). Robust statistical methods in R using the WRS2 package. Behavior Research Methods, 52(2), 464–488. 10.3758/s13428-019-01246-w

Mirman, D., Dixon, J. A., & Magnuson, J. S. (2008). Statistical and computational models of the visual world paradigm: Growth curves and individual differences. Journal of Memory and Language, 59(4), 475–494. 10.1016/j.jml.2007.11.006

Morita, T., Slaughter, V., Katayama, N., Kitazaki, M., Kakigi, R., & Itakura, S. (2012). Infant and adult perceptions of possible and impossible body movements: An eye-tracking study. Journal of Experimental Child Psychology, 113(3), 401–414. 10.1016/j.jecp.2012.07.003

Oberman, L. M., Ramachandran, V. S., & Pineda, J. A. (2008). Modulation of mu suppression in children with autism spectrum disorders in response to familiar or unfamiliar stimuli: The mirror neuron hypothesis. Neuropsychologia, 46(5), 1558–1565. 10.1016/j.neuropsychologia.2008.01.010

Pangelinan, M. M., Kagerer, F. A., Momen, B., Hatfield, B. D., & Clark, J. E. (2011). Electrocortical dynamics reflect age-related differences in movement kinematics among children and adults. Cerebral cortex (New York, N.Y. : 1991), 21(4), 737–747. 10.1093/cercor/bhq162

Papesh M. H., & Goldinger S. D. (2012). Pupil-BLAH-metry: cognitive effort in speech planning reflected by pupil dilation. Attention, perception & psychophysics, 74(4), 754–765. doi:10.3758/s13414-011-0263-y

Preuschoff, K., ‘t Hart, B. M., & Einhäuser, W. (2011). Pupil Dilation Signals Surprise: Evidence for Noradrenaline’s Role in Decision Making. Frontiers in neuroscience, 5, 115. 10.3389/fnins.2011.00115

Reid, V., Belsky, J., & Johnson, M.H. (2005). Infant perception of human Action : Toward a developmental cognitive. Cognition. Brain, Behavior, 9(3), 193–210.

Ripoll, H., Keller, J., Olivier, I. (1994). Le développement du comportement moteur de l’enfant : l’exemple des saisies et des interceptions de balle. Enfance, 2-3, 265–284; doi : 10.3406/enfan.1994.2104

Rizzolatti, G., & Fabbri-Destro, M. (2010). Mirror neurons: From discovery to autism. Experimental Brain Research. 10.1007/s00221-009-2002-3

Rizzolatti, G., Fabbri-Destro, M., & Cattaneo, L. (2009). Mirror neurons and their clinical relevance. Nature Clinical Practice Neurology. 10.1038/ncpneuro0990

Sirois, S., & Brisson, J. (2014). Pupillometry. Wiley interdisciplinary reviews. Cognitive science, 5(6), 679– 692. 10.1002/wcs.1323

Sirois, S., & Jackson, I. R. (2012). Pupil Dilation and Object Permanence in Infants. Infancy : the official journal of the International Society on Infant Studies, 17(1), 61–78. 10.1111/j.1532-7078.2011.00096.x

Stapel, J. C., Hunnius, S., & Bekkering, H. (2012). Online prediction of others’ actions: The contribution of the target object, action context and movement kinematics. Psychological Research, 76(4), 434–445. 10.1007/s00426-012-0423-2

Troje, N. F., Westhoff, C., & Lavrov, M. (2005). Person identification from biological motion: effects of structural and kinematic cues. Perception & psychophysics, 67(4), 667–675. 10.3758/bf03193523

Van Elk, M., van Schie, H. T., Hunnius, S., Vesper, C., & Bekkering, H. (2008). You’ll never crawl alone: Neurophysiological evidence for experience-dependent motor resonance in infancy. NeuroImage, 43(4), 808–814. 10.1016/j.neuroimage.2008.07.057

Wang, C. A., & Munoz, D. P. (2015). A circuit for pupil orienting responses: implications for cognitive modulation of pupil size. Current opinion in neurobiology, 33, 134–140. 10.1016/j.conb.2015.03.018

Wickham, H., Averick, M., Bryan, J., Chang, W., McGowan, L., François, R., … Yutani, H. (2019). Welcome to the Tidyverse. Journal of Open Source Software, 4(43), 1686. 10.21105/joss.01686

